# Dietary shifts in infected mosquitoes suggest a form of self-medication despite benefits in uninfected individuals

**DOI:** 10.1101/2024.12.12.628192

**Authors:** Tiago G. Zeferino, Alfonso Rojas Mora, Armelle Vallat, Jacob C. Koella

**Author notes:** Corresponding author and (TGZ).

## Abstract

Immune responses protect against infectious diseases but often incur physiological costs, such as oxidative stress. In mosquitoes, these costs may shape behaviours that help regulate oxidative balance, potentially including the consumption of nectar containing bioactive substances like prooxidants and antioxidants. We investigated whether *Anopheles gambiae* adjust their preferences for diets with such substances when they are infected with the microsporidian parasite *Vavraia culicis*. Using sugar solutions supplemented with hydrogen peroxide (a prooxidant) or ascorbic acid (an antioxidant), we assessed the feeding preferences of uninfected and infected mosquitoes at different ages and measured the effects of these diets on oxidative homeostasis, parasite load, and lifespan. Infected mosquitoes initially preferred the prooxidant diet, which reduced parasite load and extended lifespan, before shifting their preference towards the antioxidant diet as infection progressed. In contrast, uninfected mosquitoes consistently preferred the unsupplemented sugar, likely to avoid oxidative stress induced by the supplemented diets, which, surprisingly, also increased their lifespan. These results suggest a form of self-medication, despite benefits in uninfected individuals, where mosquitoes dynamically adjust their dietary choices throughout infection. We propose that such dietary strategies may be widespread among mosquitoes, helping them manage oxidative stress, with potential implications for vector–pathogen interactions and the success of biological control programmes.

## Introduction

Many animals change their behaviour when they are infected by a parasite. While this is often interpreted as manipulation by the parasite in an attempt to increase its transmission, it can also be an adaptive response of the infected animals to increase their reproductive fitness. Anti-parasitic behaviour has indeed been observed in many animals, including arthropods [1] and primates [2]. One such behaviour is self-medication, that is, the use of secondary plant chemicals or other non-nutritive substances [3], either therapeutically as a response to parasitic infection [4] or prophylactically (often in anticipation of a high parasite risk [5–8]). However, the boundaries between nutrients, medicines and toxins are flexible and are often determined solely by the ingested dose of a chemical [9].

Self-medication was already described by Aristotle in his work “Historia animalium” (384-322 B.C.) [10]. It was initially thought to require observation, learning and conscious decision-making [2,11], and thus restricted to animals with high cognitive abilities [12]. A typical example is a chimpanzee seeking medicinal plants to treat its diseases [3,13]. However, self-medication has been observed in “less intelligent mammals” [14], but also in insects (like moths [4,15], ants [5] and fruit flies [16]), indicating innate-based medication.

Mosquitoes are intriguing species to study self-medication. As vectors carrying parasites with significant implications for human health, mosquitoes feed on nectar [17–21] and thereby ingest many biologically active compounds, some of which possess anti-parasitic properties [22]. In addition, the compounds can have oxidative or antioxidative properties, thus affecting the production of reactive oxygen species (i.e., ROS, metabolic by products) and the development of oxidative stress (i.e., OS) in mosquitoes [23], both of which mediate life-history traits [24,25] and the immune response [26]. The activation of immune responses and detoxification pathways also generates ROS as a by-product, which, if not counteracted by antioxidants, leads to oxidative stress. Therefore, the protective mechanisms that shield mosquitoes from parasites come at a cost in the form of OS, increasing the selective pressure for them to manage their oxidative homeostasis.

Finally, the plants that mosquitoes feed on have a strong influence on the outcome of their infection by, for example, malaria parasites [27], probably because of a combination of the toxic secondary metabolites and the nutritional quality of the nectar. Elevated levels of ROS, which can result from ingesting certain plant compounds, enhance the immune responses against *Plasmodium* and other pathogens [28,29]. In addition to bolstering immunity, high ROS levels can also exert direct toxic effects on parasites, though they may simultaneously harm the host if not tightly regulated.

Here, we tested whether the mosquito *Anopheles gambiae* engages in therapeutic self-medication after being infected by the microsporidian *Vavraia culicis* [30] by feeding on sugar sources supplemented with biologically relevant compounds. As nectars naturally consumed by mosquitoes vary in their composition of prooxidants, such as hydrogen peroxide, and antioxidants, such as ascorbic acid, we compared a standard sugar solution with two supplemented alternatives: one containing hydrogen peroxide and the other ascorbic acid, both at concentrations typically found in nectar [31–34]. Prooxidants can enhance immune responses by increasing oxidative stress, albeit at a physiological cost, whereas antioxidants may mitigate these costs but risk impairing parasite defence mechanisms. These dual and potentially opposing effects made these compounds particularly relevant for studying self-medication in mosquitoes.

Additionally, we examined whether mosquitoes’ dietary choices were driven by changes in parasite burden over time, focusing on parasite load at day 8 post-emergence and at death, as *V. culicis* replicates rapidly before stabilizing [30]. We also evaluated whether consuming the preferred dietary elements affected oxidative stress and longevity in infected and uninfected individuals. Finally, we explored the broader implications of this behaviour, discussing its potential widespread nature, and relevance for vector–pathogen interactions and for biological control. We further considered the biological meaning of the apparent absence of measurable costs in uninfected individuals [35,36], noting that this does not necessarily imply a true absence of cost and that hidden or unmeasured trade-offs may still exist.

## Materials and Methods

### Experimental system

We used the Kisumu strain of *An. gambiae s.s* [37], which had been maintained at our standard laboratory conditions (about 600 individuals per cage, constant access to 6% sucrose solution, 26 ± 1°C, 70 ± 5% relative humidity and 12 h light/dark) for many years before the experiments.

The microsporidian *V. culicis floridensis* was provided by J.J Becnel (USDA, Gainesville, FL, USA). Originally discovered in *Aedes albopictus,* this parasite was later found to parasitize several mosquito genera, including *Aedes*, *Culex*, *Anopheles*, *Culiseta*, *Ochlerotatus* and *Orthopodomyia* [38–40]. We maintained it by alternately infecting *Ae. aegypti* and *An. gambiae* to ensure its status as a generalist parasite, as described previously [30].

*V. culicis* is an obligatory, intracellular parasite. Mosquitoes are infected when, as larvae, they ingest the spores along with their food. After several rounds of replication, the parasite produces new infectious spores that spread to gut and fat body cells. The spores are released to larval sites when infected larvae die, when adults die on the surface of water, or when eggs covered with spores are laid onto the surface of water.

### Mosquito rearing and maintenance

Freshly hatched (0-3 hours old) larvae were individually placed into 12-well culture plates, each containing 3 ml of deionized water. The larvae received Tetramin Baby® fish food every day according to their age (0.04, 0.06, 0.08, 0.16, 0.32 and 0.6 mg/larva respectively on days 0, 1, 2, 3, 4 and 5 or older [41]).

Two-day-old larvae were exposed to either 0 or 10,000 spores of *V. culicis*. Pupae were transferred to individual 50 ml falcon tubes with approximately 10 ml of deionized water [42]. Female adults were individually moved to 150 ml cups (5.5 cm Ø x 10 cm) covered with a net. The cups contained a 50 mm Ø petri dish (to prevent mosquitoes from drowning) floating on the surface of 50 ml deionized water, along with a 10 x 7 cm rectangular filter paper to maintain humidity.

### Diets

We compared three diets: 10% sucrose (referred to as sugar diet), 10 % sucrose supplemented with 8 mM of hydrogen peroxide (prooxidant diet) and 10% sucrose supplemented with 1mg/mL of ascorbic acid (vitamin c; antioxidant diet). The doses used for the different treatments correspond to concentrations found in nectar [31–34], and had been tested before the experiment to ensure they did not kill the mosquitoes. Redox cycles induced by hydrogen peroxide and ascorbic acid have been suggested as the most likely defence mechanism against a broad spectrum of bacteria and fungi in nectar [43,44].

### Preference

The preferences of infected and uninfected mosquitoes were tested 0, 4, 8, and 12 days after emergence. These time points were selected because they correspond to key stages in the infection dynamics of *V. culicis* in *An. gambiae* [30]. Thus, no spores are detectable on day 0, spores become detectable in all infected individuals by day 4, and spore load increases until day 8. Then parasite growth slows and eventually plateaus after day 12. Until the day before each assay, mosquitoes were maintained on uncoloured cotton balls soaked in a 10% sucrose solution, which was replaced daily. During the assays, the diets were compared pairwise, that is, mosquitoes were put individually into cups and given two food choices: prooxidant and sugar, antioxidant and sugar, or prooxidant and antioxidant. The solutions were coloured with pink or green food dye (Trawosa®). An equal number of mosquitoes were tested for each diet in pink and green to prevent a possible preference for the colour from biasing the results.

Each food choice was presented via a tightly compressed cotton ball (approx. 4 cm diameter) that was soaked with about 5–8 ml of the respective solution and placed onto the net of the cup. The two food sources were approximately 3 cm apart. The cottons were freshly soaked just before testing and placed onto the net at the same time to ensure equal accessibility. Each individual was tested once and the placement (left/right) of the cotton balls was randomised among cups.

Mosquitoes were starved for 12 hours before the assay to enhance feeding motivation. Pilot tests had shown that this period induced nearly all mosquitoes to feed within five minutes. They were given access to the two diets for 30 minutes. They were then placed into 2 ml Eppendorf tubes and immediately frozen at –20L°C to prevent digestion. The colour of the abdomen was assessed under a dissecting microscope to determine diet choice.

We scored mosquitoes as either green, pink, or "non-attributable" (NA) without knowing which colour corresponded to which diet. The NA category referred to mosquitoes that had consumed both diets, resulting in a mixed abdominal colour. These individuals (between 0 and 3 per treatment) were excluded from further analysis, as the mixing of dyes in the gut prevented us from determining diet preference or feeding chronology.

### Wing length and spore load

We assessed the size of each mosquito from its right wing, measuring the distance from the axillary incision to the tip [45,46] with the software ImageJ v1.54 [47]. The remaining portion of the mosquito, together with a stainless-steel bead (Ø 5 mm), was put into a 2 ml Eppendorf tube containing 0.1 ml of deionised water, and homogenised with a Qiagen TissueLyser LT at 30 hz for two minutes. 0.1 µl of each sample was added to a haemocytometer and the spores were counted at 400x magnification. All of the exposed mosquitoes from this experiment harboured spores.

### Oxidative stress

Individuals were reared as previously described until they emerged as adults. Infected and uninfected female adults then received one of the diets (uncoloured cotton balls soaked in the sugar, the prooxidant or the antioxidant solution) for seven days. The next day, alive individuals were collected in a 2 ml Eppendorf tubes, weighed, snap-frozen and then stored at -80° C. We then measured oxidative stress in all individuals, and spore load (as described above) in infected ones.

We assessed oxidative stress [48,49] with a combination of the reduced (GSH) and the oxidised (GSSH) form of glutathione using an HPLC-MS. GSH neutralizes reactive oxygen species with a chemical reaction that converts it into GSSG [50]. Our three measures of oxidative stress were therefore the amount of GSH, the amount of GSSG and, the proportion of GSSG over the total glutathione (i.e., tGSH = GSH + GSSG), which is a reliable indicator of cellular oxidative stress [51]. Note that an analysis of the total glutathione content (GSH + GSSG) would be biased by the reduced form, since the concentration of GSH is usually much higher than that of GSSG; in our experiment there was a 25-fold difference between the two.

To measure GSH and GSSG [52] we kept the samples on ice. We added a stainless-steel bead (Ø 5 mm) and 80 µl of PBS (pH = 7.4) to each sample, and homogenised it with a Qiagen TissueLyser LT at 45 Hz for four minutes. The homogenates were centrifuged at 10 000 RPM at 4° C, and 12 µl of the supernatant was transferred to a 2 ml Eppendorf tube; the rest of the homogenate was stored at -20° C for later spore load quantification, as described for the experiment on preference. We added 10 µl of formic acid (1%), 181 µl of mqH_2_O and 5 µl of glutathione reduced ethyl ester (800ng/ml, which serves as the internal standard; Sigma-Aldrich®), and vortexed the samples for 10 seconds. After centrifuging the samples for 15 min at 13 000 RPM at 4° C, 180 µl of the extract was pipetted into a syringe with a filter, and the extract was gently blown into a 250 µl conical glass insert placed inside a vial.

The samples were then injected into a HPLC-MS system for quantification (**Table. 1**). The amounts of GSH (Sigma-Aldrich®) and GSSG (Sigma-Aldrich®) were based on linear regression from ten calibration points (0.5–500 ng/mL). The liquid chromatography-mass spectrometry analyses were performed on a QTRAP 6500+ (Sciex) coupled to Acquity UPLC^®^ I-class (Waters). The column used was an Acquity UPLC^®^ HSS T3 (1.8 µm, 2.1 x 100 mm) at a temperature of 25°C. The mobile phases were milliQ water-1mM ammonium formate-0.05% formic acid (A) and acetonitrile-0.05% formic acid (B), with the following gradient elution: 100% A to 95% A in 2 min, at 4.7 min 100%B until 6.5 min, and at 6.7 min 100% A, then re-equilibration to 100% A for 4.8 min for a total run time of 11.5 minutes. The flow rate was 0.4 mL/min, with the volume injected equal to 2 µl. The source (ESI) parameters of the analysis were: ion spray voltage: 5500 V; curtain gas: 35; Ion source gas: GS1 40 and GS2 35; temperature: 550 °C.

**Table 1.** MRM transitions, declustering potential (DP), collision energy (CE), and collision cell exit potential (CXP) for GSH, GSSG, and IS detection and quantification.

| Compound | MRM transition | DP (V) | CE (V) | CXP (V) | Dwell time (msec) |
| --- | --- | --- | --- | --- | --- |
| GSH Quanti | 308.0 → 179.0 | 56 | 17 | 14 | 30 |
| GSH Quali | 308.0 → 162.0 | 66 | 33 | 14 | 30 |
| GSSG Quanti | 613.2 → 355.1 | 16 | 33 | 24 | 30 |
| GSSG Quali | 613.2 → 484.2 | 21 | 31 | 12 | 30 |
| IS | 336.0 → 190.0 | 71 | 21 | 14 | 30 |

With these settings, the limit of detection (LOD) for GSH was 0.07 ng/ml while the limit of quantification (LOQ) was 0.25 ng/ml. The LOD for GSSG was 0.5 ng/ml, and the LOQ was 2 ng/ml. The retention times were for GSH 2.06 min, GSSG 2.83 min and IS (GSH-ee) 3.60 min. Detection in ESI positive mode and for the quantification MRM (Multiple Reaction Monitoring) mode was used.

### Longevity

Individuals were reared as previously described until they emerged. Infected and uninfected female adults then received one of the diets (uncoloured cotton balls soaked in sugar, prooxidant, or antioxidant solution) for seven days. From day 8 onward, all mosquitoes were provided with unsupplemented sugar until their death. The longevity of each individual was recorded daily. At death, the spore load of infected mosquitoes was measured as previously described.

### Statistical analyses

All analyses were conducted with the R software [53] version 4.4.2, using the packages DHARMa [54], car [55], glmmTMB [56], emmeans [57] and multcomp [58]. Significance was assessed with the “Anova” function of the “car” package [55]. We used a type III ANOVA in the case of a significant interaction and a type II ANOVA otherwise. When relevant, we performed post-hoc multiple comparisons using the package “emmeans” with the default Tukey adjustment for multiple testing. To assess whether dietary choices deviated from an equal (50:50) distribution, we also used “emmeans” with the default Holm correction. When relevant, we also reported effect sizes to complement statistical significance. Effect sizes were expressed as *Cohen’s d* for Gaussian linear models, *Odds Ratios* (OR) for Binomial generalized linear models (GLMs), and *Rate Ratios* (RR) for Poisson, Gamma or Negative Binomial GLMs.

The preference for a diet of each mosquito was coded as a binary response. Preference in each choice test (prooxidant vs. sugar, antioxidant vs. sugar, prooxidant vs. antioxidant) was analysed with a generalised linear model with a binomial distribution of errors, where the explanatory variables were age, infection status and their interaction. In a preliminary analysis we had included wing length and colour of the diet, but omitted both factors from the results shown here, for they explained very little of the variance.

We also considered the impact of spore load on this preference by analysing only infected individuals with a generalised linear model with a binomial distribution of errors, in which infection status was replaced by spore load, and wing length was added as a covariable. Because spore load did not affect this preference, the results are only briefly summarised in the results section and are analysed in detail in the Supplementary Information.

The GSH content was analysed with a linear model with a Gaussian distribution of the errors, and the GSSG content was analysed with a generalised linear model with a negative binomial distribution of errors, both with infection status, diet and their interaction as explanatory variables, and mosquito weight as a covariable.

The ratio of GSSG to total glutathione (tGSH) was analysed with a generalised linear model with a Gamma distribution of errors, where infection status, diet and their interaction, were the explanatory variables and mosquito weight was a covariable.

The presence of spores at day 8 was coded as a binary response (presence or absence of detectable spores), and analysed with a generalised linear model with a binomial distribution of errors; the spore load of individuals with positive counts at day 8 was analysed with a generalised linear model with a quasi-Poisson distribution of errors. In both analyses the explanatory variables were diet, GSH content, GSSG content, and their interactions with each of these variables, and weight was included as a covariable. Additionally, the spore load at death for mosquitoes that were maintained on the sugar diet from day 8 onwards was analysed with a generalised linear model with a negative binomial distribution of errors, where infection status, diet and their interaction as explanatory variables. Because we counted the number of spores in a haemocytometer containing 0.1 µl of the sample (i.e. 1/1000 of the total volume), the detection threshold was estimated to be 1000 spores.

Survival up to day 8 was coded as a binary response (alive or dead), and analysed with a generalised linear model with a binomial distribution of errors; longevity (age at death) of mosquitoes maintained on the sugar diet from day 8 onwards was analysed with linear model with a Gaussian distribution of the errors. In both analyses, the explanatory variables where infection status, diet and their interaction.

## Results

### Preference

We assessed the dietary preferences of infected and uninfected mosquitoes at 0, 4, 8, and 12 days after adult emergence. Prior to the preference assays, all mosquitoes were maintained on uncoloured cotton balls soaked in a 10% sucrose solution, which was replaced daily. During the assays, mosquitoes (individually placed in cups) were presented with two diet choices in pairwise combinations: prooxidant vs. sugar, antioxidant vs. sugar, or prooxidant vs. antioxidant (see **Fig. 1A** for experimental design).

**Figure 1.**
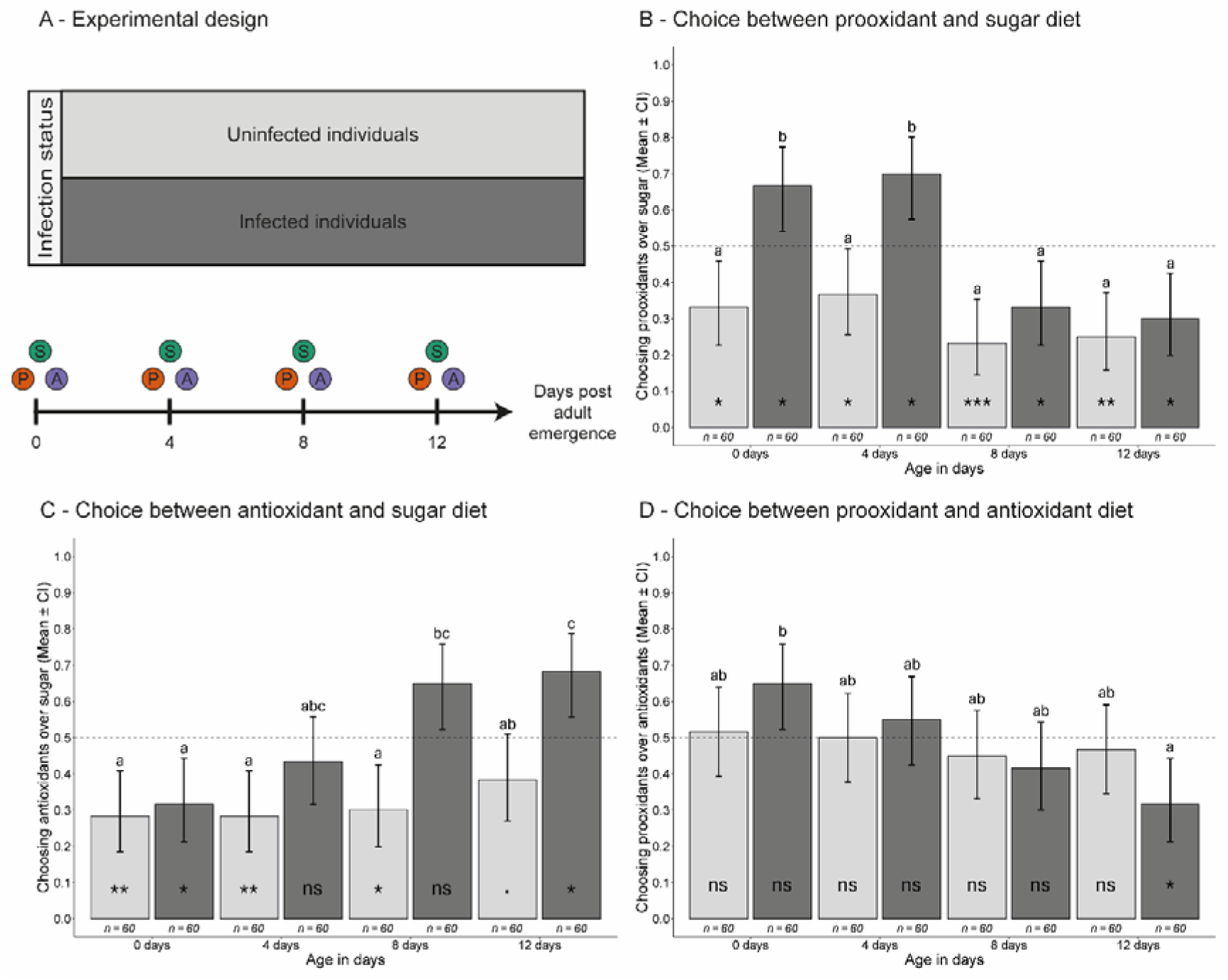
Dietary preferences of infected (dark bars) and uninfected mosquitoes (light bars) across different age classes when given two food choices. **(A)** Schematic of the experimental design. The proportion of mosquitoes **(B)** choosing the prooxidant diet when offered a choice between prooxidant and sugar diets, **(C)** choosing the antioxidant diet when offered a choice between antioxidant and sugar diets, and, **(D)** choosing the prooxidant diet when offered a choice between prooxidant and antioxidant diets. Bars represent the mean proportion of mosquitoes choosing the indicated diet, with sample sizes shown below each bar, and error bars showing the 95% confidence intervals of the means. Letters indicate statistically significant differences from multiple comparisons corrected using Tukey’s method within each choice. Asterisks inside bars indicate whether the observed preference significantly deviates from a random (50:50) choice: ns = not significant (p > 0.1),. = marginally not significant (0.1 ≥ p > 0.05), * = p ≤ 0.05, ** = p ≤ 0.01, *** = p ≤ 0.001.

#### Prooxidant vs Sugar

When given the choice between the prooxidant and sugar diets, approximately 60% chose the sugar diet, while 40% selected the prooxidant diet.

Infected individuals were significantly more likely to feed on the prooxidant diet than uninfected ones (50% vs. 29%; χ^2^ = 20.85, df = 1, p < 0.001), with an odds ratio of 2.42 (95% CI: 1.63–3.57). This preference was strongest early in infection, with 67% of infected mosquitoes choosing the prooxidant at day 0 (OR = 4.00, 95% CI: 1.87–8.55) and 70% at day 4 (OR = 4.03, 95% CI: 1.88–8.63). The effect gradually declined at later time points, reaching 33% at day 8 (OR = 1.64, 95% CI: 0.73–3.67) and 30% at day 12 (OR = 1.29, 95% CI: 0.57–2.87; χ^2^ = 27.79, df = 3, p < 0.001; **Fig. 1B**). Spore load among infected mosquitoes did not significantly influence dietary preference (χ^2^ = 0.02, df = 1, p = 0.877; see **Table S1**).

#### Antioxidant vs Sugar

When mosquitoes were offered a choice between the antioxidant and sugar diets, 58% selected sugar and 42% chose the antioxidant. Infected mosquitoes were significantly more likely to choose the antioxidant diet than uninfected ones (52% vs. 31%; χ^2^ = 20.91, df = 1, p < 0.001), with an OR of 2.42 (95% CI: 1.65–3.54). Unlike the prooxidant comparison, this preference emerged later in life, becoming evident at day 8 (65% selection; OR = 4.33, 95% CI: 2.01–9.32) and persisting at day 12 (68% selection; OR = 3.47, 95% CI: 1.63–7.37). Overall, preference for the antioxidant diet increased with age (χ^2^ = 16.44, df = 3, p < 0.001; **Fig. 1C**). Spore load did not significantly affect dietary preference among infected mosquitoes (χ^2^ = 0.18, df = 1, p = 0.671; see **Table S2**).

#### Prooxidant vs Antioxidant

When directly comparing the prooxidant and antioxidant diets, mosquitoes showed no clear preference: 48% selected the prooxidant diet, and 52% chose the antioxidant diet. Dietary choice did not significantly differ between infected and uninfected mosquitoes (both 48%; χ^2^ = 0.00, df = 1, p = 0.995), with an OR of 0.99 (95% CI: 0.69–1.43). However, dietary preference shifted with age: younger mosquitoes tended to favour the prooxidant diet, while older mosquitoes increasingly preferred the antioxidant diet (χ^2^ = 10.19, df = 3, p = 0.017; **Fig. 1D**). Importantly, unlike the previous comparisons (**Figs. 1B** and **1C**), most choices did not significantly deviate from a 50:50 distribution (as indicated by the absence of asterisks in **Fig. 1D**). Spore load did not significantly influence dietary choice (χ^2^ = 0.06, df = 1, p = 0.805; see **Table S3**).

### Oxidative stress

To assess oxidative stress levels, infected and uninfected female mosquitoes were maintained individually on one of three diets: sugar, prooxidant, or antioxidant, provided on uncoloured cotton balls soaked in the respective solutions. Mosquitoes received these diets from emergence until seven days post-emergence. The following day, all surviving individuals were collected for oxidative stress measurements (see **Fig. 2A** for experimental design).

**Figure 2.**
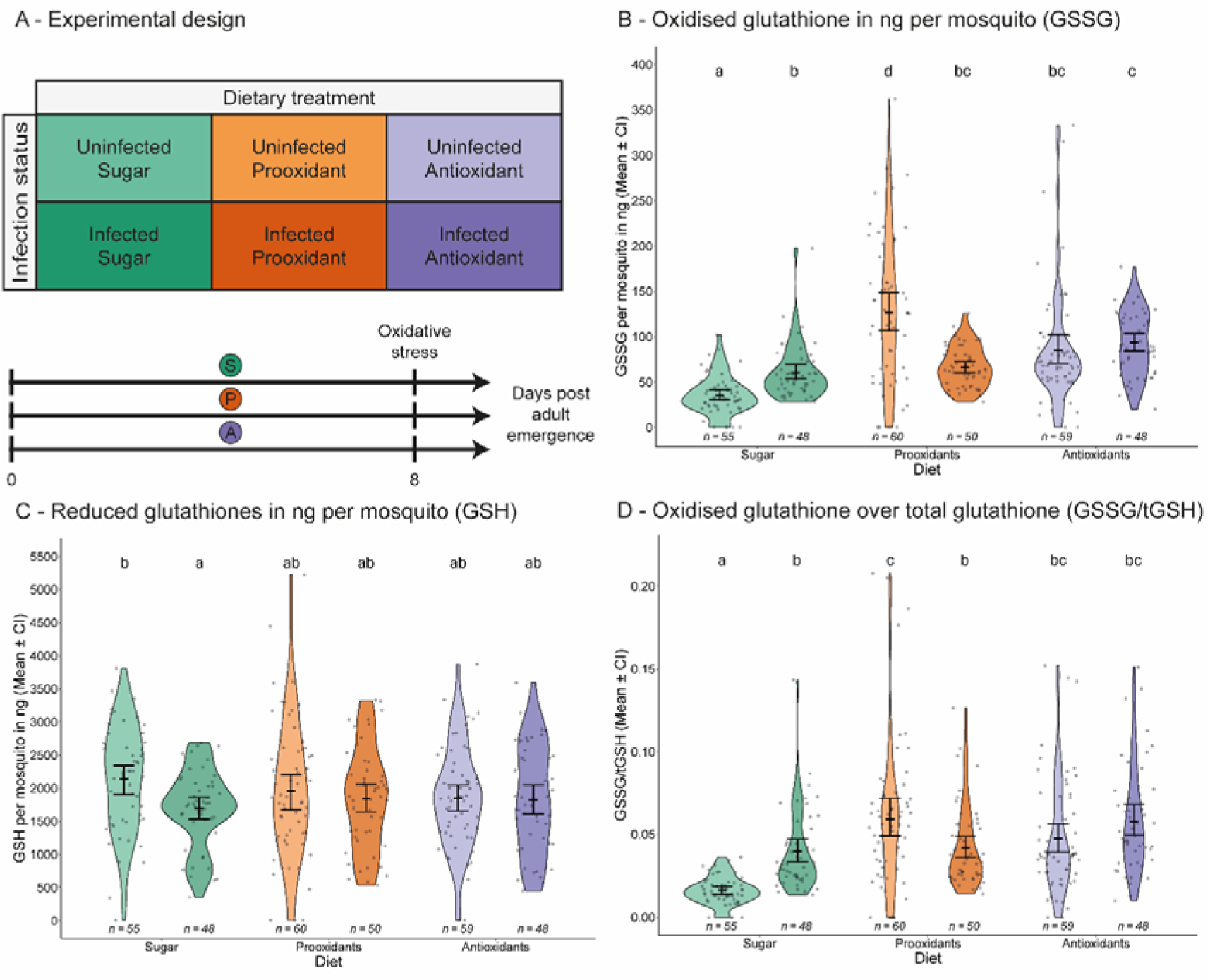
Oxidative stress of infected (dark colours) and uninfected mosquitoes (light colours) after consuming the specified diet for 7 days. **(A)** Schematic of the experimental design. **(B)** Oxidised glutathione (GSSG) and **(C)** reduced glutathione (GSH) content in ng per mosquito. Note that the scales differ between GSSG and GSH content. The proportion of **(D)** oxidised glutathione per mosquito, calculated by dividing the GSSG content over the total glutathione (i.e., tGSH = GSH + GSSG). Panels **(B–D)** show individual data points, mean values, 95% confidence intervals of the mean, and violin plots to illustrate the distribution within each treatment. Sample sizes are indicated below each treatment group. Letters indicate statistically significant differences from multiple comparisons corrected using Tukey’s method.

#### Glutathione oxidised (GSSG)

The amount of GSSG did not differ between infected and uninfected mosquitoes (72 ng vs 73 ng; χ^2^ = 0.15, df = 1, p = 0.702 with a rate ratio (RR) of 0.99 (95% CI: 0.85– 1.14). In contrast, diet strongly influenced GSSG levels. Mosquitoes fed the prooxidant diet (92 ng; RR = 2.00, 95% CI: 1.66–2.43) or the antioxidant diet (90 ng; RR = 1.94, 95% CI: 1.61–2.35) had significantly higher GSSG levels than those fed unsupplemented sugar (46 ng; χ^2^ = 70.14, df = 2, p < 0.001). There was a significant interaction between infection status and diet (χ^2^ = 41.80, df = 2, p < 0.001; **Fig. 2B**): infected mosquitoes on the sugar diet had substantially higher GSSG levels than uninfected ones (60 ng vs. 35 ng; RR = 1.70, 95% CI: 1.31–2.21), while GSSG levels of infected and uninfected mosquitoes were similar on the antioxidant diet (93 ng vs. 86 ng; RR = 1.08, 95% CI: 0.83–1.39). Interestingly, infected mosquitoes on the prooxidant diet had lower GSSG levels than uninfected ones (67 ng vs. 128 ng; RR = 0.52, 95% CI: 0.41–0.67).

#### Glutathione reduced (GSH)

In contrast, infected mosquitoes had significantly lower levels of GSH than uninfected ones (1783 ng vs 1985 ng; χ^2^ = 4.74, df = 1, p = 0.029; Cohen’s d = -0.25). This reduction was consistent across diets, as neither diet (χ^2^ = 0.63, df = 2, p = 0.730) nor the interaction between infection status and diet (χ^2^ = 3.67, df = 2, p = 0.160) significantly influenced GSH levels (**Fig. 2C**). While the interaction was not statistically significant, post hoc comparisons suggest that the difference in GSH between infected and uninfected individuals was more pronounced in sugar-fed mosquitoes (Cohen’s d = -0.54), indicating that diet may modulate the magnitude of the infection effect, even if this was not detected in the overall analysis.

#### GSSG/tGSH

If oxidative stress was assessed as the ratio between GSSG and total glutathione, infected mosquitoes exhibited had higher oxidative stress than uninfected ones (0.0464 vs. 0.0430; χ^2^ = 5.95, df = 1, p = 0.015), with a rate ratio of 1.26 (95% CI: 1.06–1.50). Diet strongly influenced oxidative stress, with higher levels in mosquitoes fed prooxidant (0.0533; RR = 2.00, 95% CI: 1.66– 2.43) or antioxidant diets (0.0520; RR = 2.00, 95% CI: 1.66–2.43) than in those fed sugar (0.0272; χ^2^ = 54.35, df = 2, p < 0.001). There was a significant interaction between diet and infection status (χ² = 35.05, df = 2, p < 0.001; **Fig. 2D**). Specifically, among sugar-fed mosquitoes, oxidative stress was higher in infected than in uninfected individuals (0.0396 vs. 0.0162; RR = 2.42, 95% CI: 1.78–3.30), but among those fed the antioxidant diet it was similar (0.0580 vs. 0.0472; RR = 1.24, 95% CI: 0.91– 1.68), and among the prooxidant-fed ones it was lower in infected than uninfected mosquitoes (0.0418 vs. 0.0629; RR = 0.66, 95% CI: 0.49–0.89). These results were consistent with post-hoc multiple comparisons and highlight diet-dependent differences in oxidative stress response.

### Spore load

Spore load was measured in two independent experiments. In both, infected and uninfected female mosquitoes were maintained on either the sugar, prooxidant, or antioxidant diets, provided on uncoloured cotton balls soaked in the respective solutions, until seven days post-emergence. In the first experiment, all surviving individuals were collected the following day for spore load quantification. In the second experiment, mosquitoes were switched to a sugar-only diet (replaced daily) from day 8 onwards and maintained individually until death, at which point their spore load was measured (see **Fig. 3A** for experimental design).

**Figure 3.**
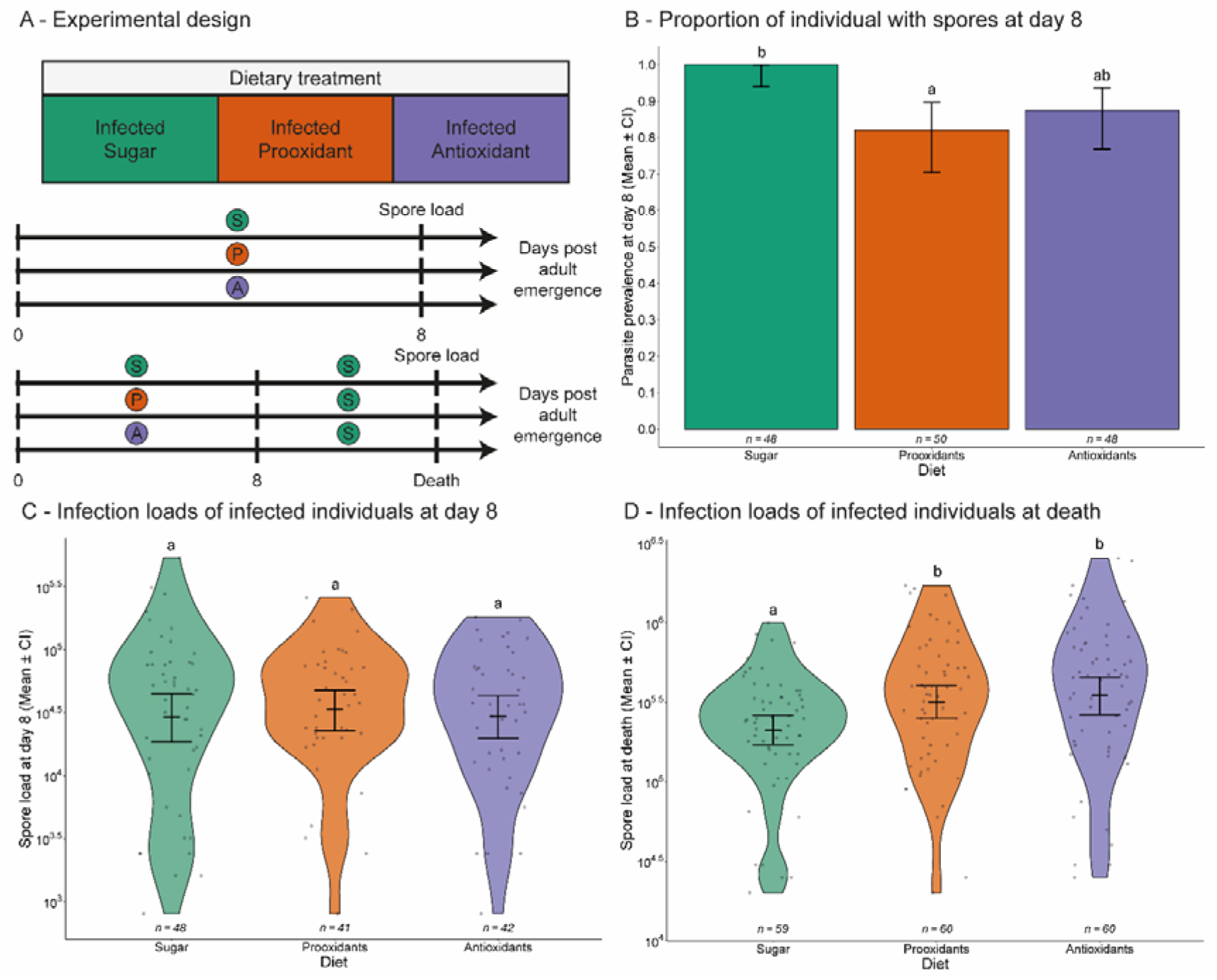
Parasite load of infected mosquitoes after consuming a specific diet for 7 days. **(A)** Schematic of the experimental design. **(B)** The proportion of mosquitoes that had detectable spores at day 8. Bars represent mean proportions, with 95% confidence intervals shown as error bars. **(C)** Spore load at day 8 or **(D)** at death, shown only for mosquitoes with positive counts. Panels **(C–D)** show individual data points, mean values, 95% confidence intervals of the mean, and violin plots to illustrate the distribution within each treatment. Sample sizes are indicated below each treatment group. Letters indicate statistically significant differences from multiple comparisons corrected using Tukey’s method.

#### Parasite prevalence at day 8

All mosquitoes fed on sugar harboured spores, whereas only 82% of those fed on prooxidant (82%) and 87% of those fed on antioxidant did (χ^2^ = 10.70, df = 2, p = 0.005; **Fig. 3B**). The likelihood of harbouring spores decreased significantly with increasing GSSG levels (χ^2^ = 5.03, df = 1, p = 0.025). In contrast, GSH levels (χ^2^ = 2.95, df = 1, p = 0.086) and the interaction terms (diet * GSH: χ^2^ = 0.68, df = 2, p = 0.712; diet * GSSG: χ^2^ = 0.06, df = 2, p = 0.970; GSH * GSSG: χ^2^ = 1.49, df = 1, p = 0.221; diet * GSH * GSSG: χ^2^ = 0.00, df = 2, p = 0.998) had no significant effects on parasite prevalence.

#### Spore load on day 8

Among infected mosquitoes, the spore load on day 8 was not significantly affected by diet (χ^2^ = 1.07, df = 2, p = 0.584; **Fig. 3C**), with mosquitoes fed on prooxidants (5.7 × 10^4^ spores; RR = 0.70, 95% CI: 0.43–1.16) or antioxidants (5.3 × 10^4^ spores; RR = 0.66, 95% CI: 0.38–1.12) having similar spore loads as those fed on sugar (8.1 × 10^4^ spores). Spore load at this stage was also independent of GSSG (χ^2^ = 0.12, df = 1, p = 0.728), GSH levels (χ^2^ = 1.42, df = 1, p = 0.234), or their interactions with diet (diet * GSH: χ^2^ = 3.89, df = 2, p = 0.143; diet * GSSG: χ^2^ = 1.35, df = 2, p = 0.509; GSH * GSSG: χ^2^ = 0.07, df = 1, p = 0.796; diet * GSH * GSSG: χ^2^ = 4.84, df = 2, p = 0.089).

#### Spore load at death

When mosquitoes were switched to a sugar-only diet from day 8 onwards, spore load at death was significantly influenced by their earlier dietary treatment. Mosquitoes that had consumed prooxidants (4.3 × 10^5^ spores; RR = 1.56, 95% CI: 1.13–2.17) or antioxidants (5.5 × 10^5^ spores; RR = 2.02, 95% CI: 1.48–2.74) for the first seven days harboured substantially more spores at death than those that consumed sugar throughout life (2.7 × 10^5^ spores; χ^2^ = 21.73, df = 2, p < 0.001; **Fig. 3D**). This effect was independent of mosquito age at death (χ^2^ = 0.02, df = 1, p = 0.886) and showed no significant interaction between age and diet (χ^2^ = 2.40, df = 2, p = 0.300).

### Longevity

Longevity was assessed through two independent experiments. In both, infected and uninfected female mosquitoes were individually maintained on one of three diets (sugar, prooxidant, or antioxidant) provided on uncoloured cotton balls soaked in the respective solutions until seven days post-emergence. In the first experiment, survival was recorded at day 8. In the second experiment, mosquitoes were switched to a sugar-only diet from day 8 onwards (replaced daily), and their age at death was recorded (see **Fig. 4A** for experimental design).

**Figure 4.**
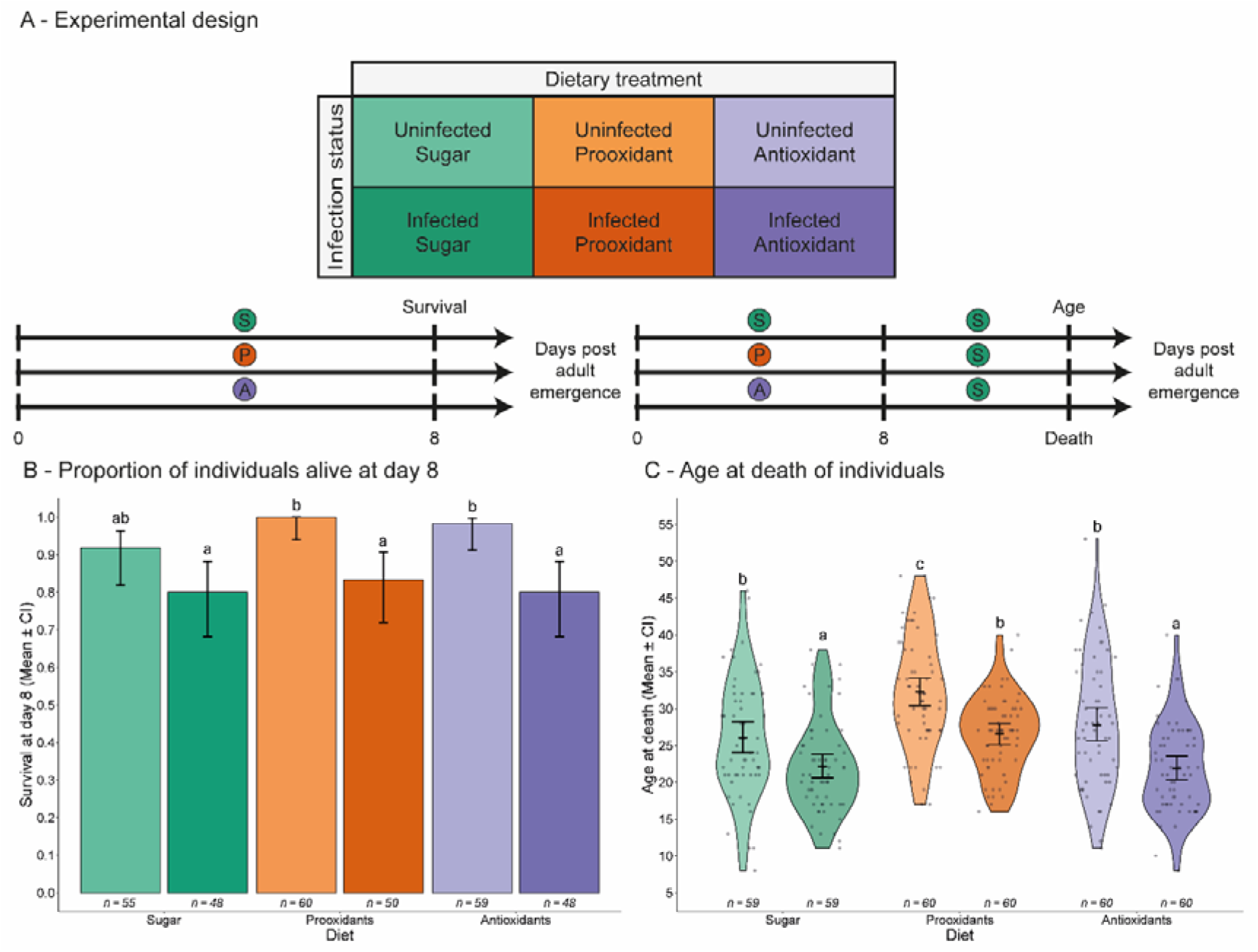
Survival at day 8 and longevity of infected and uninfected mosquitoes after consuming a specific diet for 7 days. **(A)** Schematic of the experimental design. **(B)** The proportion of mosquitoes alive at day 8. Bars represent mean proportions, with 95% confidence intervals shown as error bars. **(C)** Age at death, displayed as individual data points, mean values, 95% confidence intervals of the mean, and violin plots to illustrate the distribution within each treatment. Sample sizes are indicated below each treatment group. Letters indicate statistically significant differences from multiple comparisons corrected using Tukey’s method.

#### Survival at day 8

Overall, 88% of the mosquitoes survived until day 8, but survival was significantly lower for infected mosquitoes than for uninfected ones (81% vs 97%; χ^2^ = 7.68, df = 1, p = 0.005, **Fig. 4B**). Dietary treatment had no significant effect on survival (sugar: 85%, prooxidant: 92%, antioxidant: 89%; χ^2^ = 0.62, df = 2, p = 0.734), nor was there a significant interaction between infection status and diet (χ^2^ = 1.95, df = 2, p = 0.376).

#### Longevity

On average, mosquitoes lived 26.1 days, but infected mosquitoes had significantly shorter lifespans than uninfected ones (23.5 days vs 28.7 days; χ^2^ = 48.17, df = 1, p < 0.001; Cohen’s d = - 0.72). Diet had a significant effect on lifespan (χ^2^ = 39.17, df = 2, p < 0.001), with mosquitoes fed prooxidants living longer than those fed sugar (29.4 days vs 24.1 days; Cohen’s d = 0.73. There was no significant interaction between infection status and diet (χ^2^ = 1.46, df = 2, p = 0.481; **Fig. 4C**). This resulted in prooxidants extending lifespan in both infected (26.5 days vs 22.2 days; Cohen’s d = 0.86) and uninfected mosquitoes (32.2 days vs 26.1 days; Cohen’s d = 0.61) compared to sugar-fed mosquitoes.

## Discussion

Infection by *V. culicis* induced large shifts in mosquito dietary preferences, and maintaining mosquitoes on these diets for seven days disrupted oxidative homeostasis, delayed parasite development, and increased longevity. While this suggests an adaptive value to the dietary shifts and thus a form of self-medication, it must be noted that uninfected mosquitoes lived longer if they were fed the prooxidant diet, even if they avoided it when they had a choice. In the following sections, we first explore how these dietary choices may broadly influence mosquitoes’ oxidative balance, potentially in response to various stressors. We then discuss the dynamic, time-dependent dietary shifts during *V. culicis* infection and their consequences for parasite development and mosquito longevity. We briefly consider the implications for the use of *V. culicis* in biological control programmes. Finally, we reflect on the absence of an apparent cost in uninfected mosquitoes, discussing whether this challenges the current classification criteria for self-medication [6,59], and outline promising avenues for future research, including potential transgenerational effects.

Hydrogen peroxide and ascorbic acid, both naturally present in wild nectar sources [31–34], play critical roles as natural defences against a variety of bacteria and fungi [43,44]. We propose that their consumption by mosquitoes may reflect a behavioural response to infection-induced physiological changes—such as elevated oxidative stress—rather than a direct detection of infection itself. These shifts may occur independently of any adaptive intent, and are consistent with the known protective effects of these compounds against other stressors such as insecticides [60]. Such behaviours may be more widespread than previously recognised and could serve as a strategy for mosquitoes to manage the oxidative imbalance, not only during infection but also in response to other oxidative challenges such as blood meal ingestion [61,62] and digestion [29,63], which are essential for reproduction. Although this suggests that mosquitoes might be adapted to dealing with fluctuating oxidative environments, empirical evidence for ecologically relevant concentrations of pro- and antioxidants in field-accessible nectar sources remains limited and further field validation is needed. However, should future field studies confirm these dietary responses, it is plausible that such behaviour may not be specific to *V. culicis* infections. *An. gambiae* might exhibit similar dietary responses to other pathogens, including *Plasmodium falciparum*. This remains an open and intriguing question that warrants future investigation, particularly given the fundamental differences in infection dynamics, tissue tropism, and the duration of the extrinsic incubation period between *Plasmodium* and *Vavraia*. Understanding whether mosquitoes can adaptively adjust their diet in response to *Plasmodium* infection could have important implications for malaria transmission and its epidemiology.

In our study, infection altered mosquitoes’ dietary preferences away from the sugar diet, but the parasite burden did not influence these choices, suggesting that mosquitoes responded qualitatively rather than quantitatively to infection. Younger infected individuals preferred the prooxidant diet (**Fig. 1B**), but as they aged, their preference gradually shifted towards the antioxidant diet (**Fig. 1C**). This divergence from the sugar diet could represent an adaptive response to infection, potentially helping mosquitoes mitigate the physiological costs of harbouring the parasite. Supporting this, our oxidative stress markers show that infection induced a two-fold increase in oxidative stress (**Fig. 2D**) and reduced lifespan by approximately four days when comparing uninfected and infected individuals on the sugar diet (26.1 days vs. 22.2 days, **Fig. 4C**). Interestingly, oxidative stress levels in infected mosquitoes remained comparable across all diets, suggesting that the prooxidant and antioxidant diets were not more physiologically costly than the sugar diet, at least in terms of oxidative balance. Crucially, consuming prooxidants for seven days reduced parasite prevalence by 18% at day 8 (**Fig. 3B**) and increased the lifespan of infected mosquitoes by an average of four days (26.5 days vs. 22.2 days, **Fig. 4C**), effectively restoring their longevity to levels similar to uninfected individuals on the sugar diet (26.5 days vs. 26.1 days). This finding is consistent with a form of therapeutic self-medication, where infected animals selectively consume substances that reduce pathogen load or mitigate the costs of infection, even when these substances may be harmful in other contexts [4,7,15]. While these results suggest potential adaptive value, we acknowledge that confirming this would require measuring reproductive success, which was beyond the scope of this study.

The infection dynamics of *V. culicis* in *An. gambiae* are well described in the literature [30], with sugar-fed mosquitoes typically reaching 100% infection prevalence by day 4 post-emergence. In our study, we measured parasite prevalence slightly later, at day 8, by which point sugar-fed mosquitoes had also reached 100% prevalence, consistent with established patterns. Interestingly, mosquitoes fed prooxidants exhibited a significantly lower prevalence (82%) at this time point, suggesting that the diet may have slowed parasite proliferation. Given that full prevalence normally occurs by day 4 under sugar-only diets, it is likely that the impact of prooxidants would have been even more pronounced had we assessed parasite load earlier in the infection process. This interpretation is further supported by our newly included spore load at death data. Mosquitoes that were initially maintained on prooxidant or antioxidant diets during the first seven days, but subsequently switched back to sugar, ultimately carried higher spore loads at death compared to individuals continuously fed sugar (**Fig. 3D**). This pattern suggests that the early dietary exposure may have delayed parasite growth rather than completely suppressing it, and that reverting to sugar after day 8 permitted accelerated parasite proliferation later in life. Such a dynamic indicates a potential trade-off between early parasite suppression and longer-term tolerance [64–67], which may reflect a flexible, time-dependent dietary strategy. Alternatively, parasites may modulate host behaviour early in infection to prolong host survival and enhance transmission at later stages, as suggested by the higher parasite load observed. Thus, parasite manipulation cannot be fully excluded based on the current data, and further research will be needed to disentangle these mechanisms.

Our results further highlight a dynamic behaviour that appears to track the progression of the infection and the parasite’s growth dynamics. To our knowledge, this is the first report of such adaptive dietary changes unfolding across the course of infection. Independently of whether this response reflects a host-driven behavioural choice or parasite manipulation, mosquitoes initially favoured the prooxidant diet during the parasite’s exponential growth phase (prior to day 8, [30]), which likely hindered parasite replication at that stage (**Fig. 3B**). As the infection progressed into its slower growth phase (from day 8 onwards, [30]), mosquitoes shifted their preference towards the antioxidant diet, possibly to enhance their ability to tolerate the infection and mitigate its associated physiological costs. This smooth transition, particularly evident in 4- and 8-day-old mosquitoes (**Fig. 1D**), implies that individual conditions may mediate how these diets influence tolerance [68]. Such findings underscore the complexity of dietary choices during infection, driven by both the parasite’s life cycle and the mosquito’s physiological state. Altogether, this time-dependent pattern suggests that mosquitoes may strategically adjust their dietary choices to balance the competing demands of resistance and tolerance at different stages of infection [64–67]. These adaptive shifts in dietary preference could also have important implications for the use of microsporidians like *V. culicis* in biological control programmes. Although *V. culicis* has long been proposed as a tool to slow [69] or impede [70,71] malaria transmission, our findings indicate that mosquitoes may mitigate the fitness costs of infection through self-medication [30,71–73], potentially reducing the long-term efficacy of such control strategies.

Uninfected mosquitoes, on the other hand, consistently preferred the sugar-only diet, actively avoiding both the prooxidant (**Fig. 1B**) and antioxidant (**Fig. 1C**) diets, while showing random preferences when the sugar diet was unavailable (**Fig. 1D**). This avoidance may be linked to the oxidative stress induced by these diets, as our data show a three-fold increase in oxidative stress markers (GSSG/tGSH ratio) with antioxidants and a four-fold increase with prooxidants (**Fig. 2D**). Elevated oxidative stress is commonly considered a physiological cost, particularly as part of the immune response, which involves a trade-off between producing sufficient reactive oxygen species (ROS) to control pathogens and avoiding self-inflicted oxidative damage [26,28]. In uninfected mosquitoes, where immune challenges should be lower, the oxidative cost of these diets may be more directly harmful, as they do not appear to gain the potential protective benefits that ROS can provide in infected individuals. Interestingly, we observed that consuming prooxidants for seven days increased the lifespan of uninfected mosquitoes by an average of six days (32.2 days vs. 26.1 days, **Fig. 4C**).

One possible explanation is that moderate elevation of ROS may trigger hormetic effects, where low-level or transient stressors can enhance physiological function and longevity. This phenomenon, extensively described in the literature [74], suggests that mild oxidative stress may upregulate cellular defence mechanisms and promote overall fitness, even in absence of infection [75].

Alternatively, the absence of an apparent cost in uninfected individuals may reflect the limitations of measuring a single fitness trait (longevity) in this study. Fitness costs can be trait-specific, context-dependent, or may only emerge under certain ecological or physiological conditions. As such, the absence of a measured cost does not necessarily imply its true absence. This raises a broader conceptual question of whether the presence of a measurable cost in uninfected individuals is a strict requirement [6,59] for a behaviour to be classified as self-medication. Given the difficulty of fully excluding hidden or unmeasured costs [35,36], it may be more informative to view self-medication as a behavioural strategy that should not rely solely on the presence or absence of a cost. Instead, its classification might better rest on its functional role in mitigating the negative physiological effects of infection, as well as its avoidance by uninfected individuals, even when no apparent cost is detected. These hypotheses offer a promising avenue for further investigation and may help explain the longevity benefits we observed. Future studies could also explore whether the F1 generation from infected parents exhibits prophylactic self-medication behaviours, even in the absence of infection, or whether they adopt counter-medication strategies, such as preferring antioxidants if their parents consumed prooxidants. Such patterns would provide valuable insights into potential transgenerational effects, highlighting transgenerational costs and whether offspring can modulate their dietary choices to mitigate residual oxidative stress inherited from parental self-medication.

## Conclusion

Our study shows that *Anopheles gambiae* mosquitoes dynamically adjust their dietary preferences in response to *Vavraia culicis* infection, consistent with a form of self-medication. However, confirming the adaptive value of this behaviour would require measurements of reproductive success, which were beyond the scope of this study. Infected mosquitoes initially prefer prooxidants that reduced parasite load and extended lifespan, before shifting towards antioxidants as infection progressed. This dynamic strategy suggests a trade-off between resistance and tolerance over time. Interestingly, prooxidant consumption also prolonged the lifespan of uninfected mosquitoes, raising questions about potential hormetic effects or hidden costs. Together, these findings challenge the assumption that self-medication requires measurable costs in uninfected individuals and suggest that such behaviours could be more widespread, with important implications for mosquito–pathogen interactions and the efficacy of biological control strategies.

## Supporting information

Statistical tables

## Acknowledgements

We thank Luís M. Silva for his advice and technical support. TGZ and the project were supported by SNF grant 310030_192786.

## Author contributions

ARM conceived the overall idea. TGZ, ARM and JCK designed the experiments. TGZ collected, analysed, and interpreted the data and wrote the first draft of the manuscript. TGZ and AV developed the method to analyse the GSSG/tGSH ratio, and AV performed the sample analyses. All authors contributed critically to the drafts.

## Data availability

All data generated or analysed during this study are included as Supplementary Information files.

## Additional Information

The authors declare no competing interests.

## Notes

### Competing Interest Statement

The authors have declared no competing interest.

### Summary of Updates

This manuscript has been heavily revised and updated since the original preprint version to incorporate new data, analyses, and improvements based on feedback.

