## Supplementary material for "Dietary shifts in infected mosquitoes suggest a form of self-medication despite benefits in uninfected individuals": Statistical tables

**Table S1.** Prooxidant vs Control.

**Table S2.** Antioxidant vs Control.

**Table S3.** Prooxidant vs Antioxidant.

**Table S4.** Oxidative stress.

**Table S5.** Parasite load.

**Table S6.** Longevity

**Table S1. Prooxidant vs Control.**

| **Tested effect** |  |  |  |
| --- | --- | --- | --- |
| **Model 1a** – Preference for prooxidant | **df** | **χ^2^** | ***p*** |
| Age | 3 | 27.79 | **<0.001** |
| Infection status | 1 | 20.85 | **<0.001** |
| Age:Infection status | 3 | 6.66 | 0.084 |
| **Model 1b** – Preference for prooxidant (infected) | **df** | **χ^2^** | ***p*** |
| Age | 3 | 24.52 | **<0.001** |
| Spore load | 1 | 0.02 | 0.877 |
| Age:Spore load | 3 | 2.49 | 0.477 |

**Table S2. Antioxidant vs Control.**

| **Tested effect** |  |  |  |
| --- | --- | --- | --- |
| **Model 2a** – Preference for antioxidant | **df** | **χ^2^** | ***p*** |
| Age | 3 | 16.44 | **<0.001** |
| Infection status | 1 | 20.91 | **<0.001** |
| Age:Infection status | 3 | 6.74 | 0.080 |
| **Model 2b** – Preference for antioxidants (infected) | **df** | **χ^2^** | ***p*** |
| Age | 3 | 17.52 | **<0.001** |
| Spore load | 1 | 0.18 | 0.671 |
| Age:Spore load | 3 | 2.79 | 0.425 |

**Table S3. Prooxidant vs Antioxidant.**

| **Tested effect** |  |  |  |
| --- | --- | --- | --- |
| **Model 3a** – Preference for prooxidant | **df** | **χ^2^** | ***p*** |
| Age | 3 | 10.19 | **0.017** |
| Infection status | 1 | 0.00 | 0.995 |
| Age:Infection status | 3 | 5.42 | 0.143 |
| **Model 3b** – Preference for prooxidant (infected) | **df** | **χ^2^** | ***p*** |
| Age | 3 | 8.58 | **0.035** |
| Spore load | 1 | 0.06 | 0.805 |
| Age:Spore load | 3 | 1.54 | 0.672 |

**Table S4. Oxidative stress.**

| **Tested effect** |  |  |  |  |
| --- | --- | --- | --- | --- |
| **Model 4a –** Total GSSG |  | **df** | **χ^2^** | ***p*** |
| Infection status |  | 1 | 0.15 | 0.702 |
| Diet |  | 2 | 70.14 | **<0.001** |
| Mosquito weight |  | 1 | 0.45 | 0.504 |
| Infection status:Diet |  | 2 | 41.80 | **<0.001** |
| **Model 4b –** Total GSH |  | **df** | **χ^2^** | ***p*** |
| Infection status |  | 1 | 4.74 | **0.029** |
| Diet |  | 2 | 0.63 | 0.730 |
| Mosquito weight |  | 1 | 0.43 | 0.510 |
| Infection status:Diet |  | 2 | 3.67 | 0.160 |
| **Model 4c –** GSSG/tGSH |  | **df** | **χ^2^** | ***p*** |
| Infection status |  | 1 | 5.95 | **0.015** |
| Diet |  | 2 | 54.35 | **<0.001** |
| Mosquito weight |  | 1 | 0.18 | 0.669 |
| Infection status:Diet |  | 2 | 35.05 | **<0.001** |

**Table S5. Parasite load.**

| **Tested effect** |  |  |  |  |
| --- | --- | --- | --- | --- |
| **Model 5a** – Presence of spores at day 8 |  | **df** | **χ^2^** | ***p*** |
| Diet |  | 2 | 10.70 | **0.005** |
| Total GSH |  | 1 | 2.95 | 0.086 |
| Total GSSG |  | 1 | 5.03 | **0.025** |
| Mosquito weight |  | 1 | 0.24 | 0.623 |
| Diet:Total GSH |  | 2 | 0.68 | 0.712 |
| Diet:Total GSSG |  | 2 | 0.06 | 0.970 |
| Total GSH:Total GSSG |  | 1 | 1.49 | 0.221 |
| Diet:Total GSH:Total GSSG |  | 2 | 0.00 | 0.998 |
| **Model 5b –** Spore load at day 8 |  | **df** | **χ^2^** | ***p*** |
| Diet |  | 2 | 1.07 | 0.584 |
| Total GSH |  | 1 | 1.42 | 0.234 |
| Total GSSG |  | 1 | 0.12 | 0.728 |
| Mosquito weight |  | 1 | 0.04 | 0.833 |
| Diet:Total GSH |  | 2 | 3.89 | 0.143 |
| Diet:Total GSSG |  | 2 | 1.35 | 0.509 |
| Total GSH:Total GSSG |  | 1 | 0.07 | 0.796 |
| Diet:Total GSH:Total GSSG |  | 2 | 4.84 | 0.089 |
| **Model 5c –** Spore load at death |  | **df** | **χ^2^** | ***p*** |
| Diet |  | 2 | 21.73 | **<0.001** |
| Age at death |  | 1 | 0.02 | 0.886 |
| Diet:Total GSH:Total GSSG |  | 2 | 2.40 | 0.300 |

**Table S6. Longevity**

| **Tested effect** |  |  |  |  |
| --- | --- | --- | --- | --- |
| **Model 6a –** Survival at day 8 |  | **df** | **χ^2^** | ***p*** |
| Infection status |  | 1 | 7.68 | **0.005** |
| Diet |  | 2 | 0.62 | 0.734 |
| Infection status:Diet |  | 2 | 1.95 | 0.376 |
| **Model 6b –** Age at death |  | **df** | **χ^2^** | ***p*** |
| Infection status |  | 1 | 48.17 | **<0.001** |
| Diet |  | 2 | 39.17 | **<0.001** |
| Infection status:Diet |  | 2 | 1.46 | 0.481 |
